# Chemogenetic silencing reveals presynaptic Gi/o protein-mediated inhibition of synchronized activity in the developing hippocampus *in vivo*

**DOI:** 10.1101/2024.02.02.578550

**Authors:** Jürgen Graf, Arash Samiee, Tom Flossmann, Knut Holthoff, Knut Kirmse

## Abstract

The development of neuronal circuits is an activity-dependent process. Research aimed at deciphering the learning rules governing these developmental refinements is increasingly utilizing a class of chemogenetic tools that employ G_i/o_ protein signaling (G_i_-DREADDs) for neuronal silencing. However, their mechanisms of action and inhibitory efficacy in immature neurons with incompletely developed G_i/o_ signaling are poorly understood. Here, we analyze the impact of G_i/o_ signaling on cellular and network excitability in the neonatal hippocampus by expressing the G_i_-DREADD hM4Di in telencephalic glutamatergic neurons of neonatal mice of both sexes. Using acousto-optic two-photon Ca^2+^ imaging, we report that activation of hM4Di leads to a complete arrest of spontaneous synchronized activity in CA1 *in vitro*. Electrophysiological analyses demonstrate that hM4Di-mediated silencing is not accounted for by changes in intrinsic excitability of CA1 pyramidal cells (PCs). Instead, activation of hM4Di robustly restrains synaptic glutamate release by the first postnatal week, effectively reducing recurrent excitation in CA1. *In vivo*, inhibition through hM4Di potently suppresses early sharp waves (eSPWs) and discontinuous oscillatory network activity in CA1 of head-fixed mice before eye opening. In summary, hM4Di dampens glutamatergic neurotransmission through a presynaptic mechanism sufficient to terminate spontaneous synchronized activity in the neonatal CA1. Our findings have implications for designing and interpreting experiments utilizing G_i_-DREADDs in immature neurons and further point to a potential role of G_i/o_-dependent endogenous neuromodulators in activity-dependent hippocampal development.

**Significance Statement:** Recent advances in understanding how early activity shapes developing brain circuits increasingly rely on G_i/o_-dependent inhibitory chemogenetic tools (G_i_-DREADDs). However, their single-cellular mechanisms and efficacy in neurons with immature G_i/o_ signaling are elusive. To close this gap, we analyzed the impact of the G_i_-DREADD hM4Di in the neonatal murine hippocampus. We report that hM4Di does not mediate hyperpolarization or shunting but rather acts as a potent presynaptic silencer of glutamatergic neurotransmission in CA1 pyramidal cells. Presynaptic inhibition of excitation suffices to arrest spontaneous synchronized network activity *in vitro* and *in vivo*. Our findings provide novel insights into the generative mechanisms of spontaneous synchronized activity and bear relevance for applying chemogenetic silencing at early stages of brain development.

## Introduction

The immature rodent hippocampus displays spontaneous synchronized activity prior to the developmental onset of spatial memory and environmental exploration. In CA1 during the first and second postnatal week, activity is characterized by intermittent network bursts that alternate with periods of low activity. Network bursts are driven by input from the sensory periphery (Mohns and Blumberg, 2010; Valeeva et al., 2019; Chen et al., 2023; Gainutdinov et al., 2023; Kostka and Hanganu-Opatz, 2023), but they can also be internally generated within the entorhinal-hippocampal formation (Ben-Ari et al., 1989; Leinekugel et al., 2002; Graf et al., 2021). By co-activating large numbers of neurons (Dard et al., 2022; Graf et al., 2022), network bursts were proposed to inform nascent CA1 circuits about the fundamental statistical properties of the external (environment) and internal (body) world, thereby preparing them for later cognitive and sensory demands (for review see Cossart and Khazipov, 2022; Kirmse and Zhang, 2022).

To elucidate the learning rules underlying activity-dependent refinements in brain development, research is increasingly utilizing chemogenetic strategies, as they allow neuronal activity to be manipulated in a temporally controlled and cell type-specific manner (Donato et al., 2017; Wong et al., 2018; Murata and Colonnese, 2020; Leighton et al., 2021; Maldonado et al., 2021; Chen et al., 2023; Kostka and Hanganu-Opatz, 2023; Leprince et al., 2023). The most widely used inhibitory chemogenetic actuator is hM4Di, which is based on the human muscarinic M4 receptor employing G_i/o_ signaling for neuronal silencing (Armbruster et al., 2007). In mature neurons, endogenous neuromodulators activating G_i/o_ signaling reduce excitability by multiple mechanisms. In somatodendritic compartments, G_i/o_ mediates hyperpolarization due to activation of G-protein-coupled inwardly rectifying K^+^ (GIRK/Kir3) channels (Reuveny et al., 1994) and inhibition of Ca_v_1 channels involved in dendritic spike generation (Perez-Garci et al., 2006; Perez-Garci et al., 2013). In addition, G_i/o_-βγ dimers inhibit evoked and spontaneous vesicle release by interacting with SNARE proteins and Ca_v_2 channels in presynaptic terminals (Herlitze et al., 1996; Sakaba and Neher, 2003; Zurawski et al., 2019; Alten et al., 2022). In a manner analogous to that of endogenous G_i/o_-protein-coupled receptors, G_i_-DREADDs are known to effectively silence neuronal activity via both hyperpolarization and suppression of neurotransmitter release in the adult brain (Stachniak et al., 2014; Vardy et al., 2015). This, however, might fundamentally differ from the situation in the neonatal neocortex and hippocampus, when GIRK channel expression is low (Chen et al., 1997; Ehrengruber et al., 1997; Bony et al., 2013) and the G_i/o_-dependent hyperpolarization (Luhmann and Prince, 1991; Nurse and Lacaille, 1999; Verheugen et al., 1999) and inhibition of voltage-gated Ca^2+^ channels (Gaiarsa et al., 1995) are not yet operative.

Here, we address this issue by analyzing the single-neuron mechanisms and circuit-level efficacy of hM4Di-mediated G_i/o_-dependent silencing in the hippocampal CA1 region of developing mice *in vitro* and *in vivo*. We focus on the first and second postnatal weeks, when network activity comprises intermittent bursts that are generated in a relatively all-or-none manner (Prida and Sanchez-Andres, 1999; Flossmann et al., 2019), rendering inhibition particularly challenging. We demonstrate that the activation of hM4Di-induced G_i/o_ signaling can arrest spontaneous neuronal synchrony through the presynaptic silencing of glutamatergic neurotransmission.

## Results

### G_i/o_ protein-mediated silencing of synchronized network activity in vitro

To examine the efficacy of G_i/o_ protein-coupled chemogenetic silencing during development, we conditionally expressed hM4Di in telencephalic glutamatergic neurons from embryonic stages onward (*Emx1^IREScre^::hM4Di^LSL^* or *hM4Di^Emx1^* mice) (Gorski et al., 2002; Zhu et al., 2016; Graf et al., 2021). Of note, both CA1 pyramidal cells (PCs) and the vast majority of glutamatergic synapses in CA1 are derived from the *Emx1* lineage (Sando et al., 2017; Leprince et al., 2023). We first used three-dimensional acousto-optic two-photon Ca^2+^ imaging in CA1 *stratum pyramidale* of acute brain slices to quantify network activity at single-cell resolution at postnatal days (P) 2–5 (Fig. 1A-B). Under control conditions, spontaneous activity in CA1 PCs consisted of discrete network events, known as giant depolarizing potentials (G DPs) (Ben-Ari et al., 1989), that alternated with extended periods of low activity (Fig. 1C). Activation of hM4Di^Emx1^ by the DREADD agonist C21 (1 µM) led to a profound reduction in the mean frequency of somatic Ca^2+^ transients (CaTs) from 1.4 ± 0.4 min^−1^ to 0.0 ± 0.0 min^−1^ (n = 5 slices; Fig. 1B-E). To control for potential DREADD-independent effects of C21, we repeated these experiments in wild-type (WT) mice lacking *hM4Di^Emx1^* (n = 6 slices; Fig. 1D-E). Generalized linear mixed-effects modeling (GLMM) revealed a significant interaction term, indicating that C21 effects in slices obtained from *hM4Di^Emx1^* mice were larger than in those from WT neonates (genotype × treatment: P = 3.5 × 10^−6^; for detailed statistics, see Extended Data Table 1-1). *Post-hoc* testing confirmed that the suppression of neuronal activity by C21 was restricted to *hM4Di^Emx1^* mice, whereas C21 was ineffective in WT pups (*hM4Di^Emx1^*: P = 3.2 × 10^−3^, WT: P = 0.16, simple contrasts; Fig. 1E). Likewise, at the network level, C21-induced inhibition abolished GDPs in slices from *hM4Di^Emx1^* but not from WT neonates (Fig. 1F). Collectively,these data demonstrate effective network silencing through hM4Di^Emx1^ and identify G_i/o_ signaling as a potential inhibitory actuator of synchronized network activity in the developing hippocampus *in vitro*.

**Figure 1.**
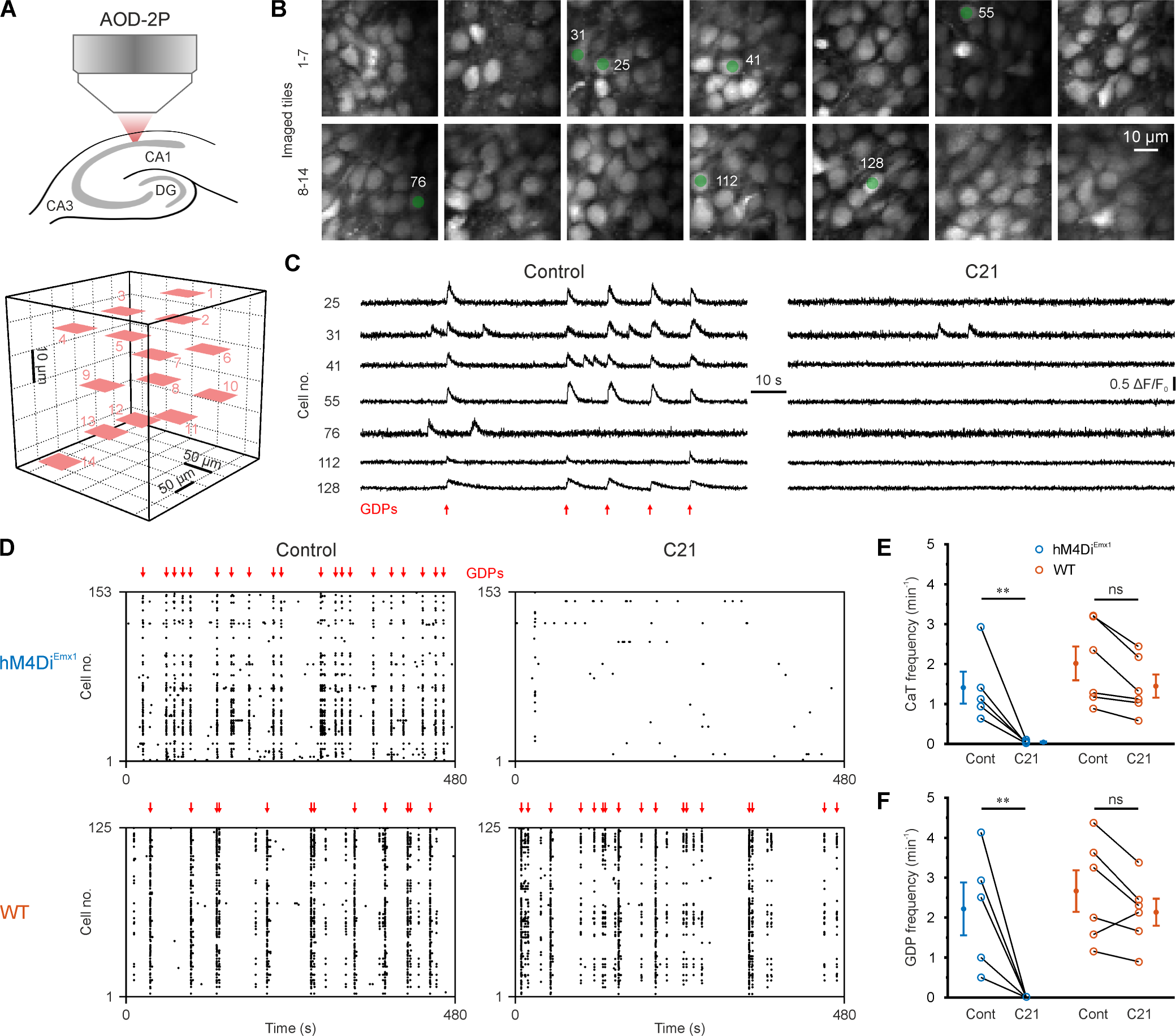
*hM4Di^Emx1^* activation profoundly inhibits synchronized network activity in the neonatal CA1 *in vitro*. ***A,*** Experimental design. *Top*: Three-dimensional (3D) Ca^2+^ imaging using an acousto-optic deflection laser-scanning two-photon microscope (AOD-2P) was performed in *stratum pyramidale* of CA1 in acute brain slices at P2–5. *Bottom*: Imaged tiles (each 50 × 50 µm) in 3D space (sample as in ***B***). ***B,*** 2P images (OGB1) of CA1 PCs (z-scored). ***C,*** Sample traces (Δ*F*⁄*F*_0_ (*t*)) of cells indicated in ***B*** before (*left*) and after (*right*) wash-in of C21 (1 µM). The slice was obtained from an *hM4Di^Emx1^* mouse. Note that GDPs (red arrows) were completely abolished by C21. ***D,*** Sample rasterplots before (*left*) and after (*right*) wash-in of C21 (1 µM) obtained from an *hM4Di^Emx1^* (*top*) and a wild-type (WT, *bottom*) mouse, respectively. Each dot represents a CaT onset in an individual cell. ***E-F***, Mean CaT frequency (***E***) and GDP frequency (***F***). Each open symbol represents a single slice. ***E-F***, Population data (closed symbols) are presented as mean ± SEM. ** P < 0.01, ns – not significant (simple contrasts). For statistics, see Extended Data Table 1-1.

### hM4Di^Emx1^-mediated silencing is not accounted for by changes in intrinsic excitability

G_i/o_ protein-coupled chemogenetic actuators are commonly used tools for neuronal silencing in the adult, as they can reduce spike rates through hyperpolarization and shunting, causing subtractive and divisive inhibition (Roth, 2016). However, these mechanisms may be less effective in the neonatal brain, as the coupling of somatodendritic GIRK channels to G_i/o_ signaling is absent or weak during the perinatal period (Fukuda et al., 1993; Chen et al., 1997; Nurse and Lacaille, 1999; Bony et al., 2013). We therefore set out to examine the mechanism(s) of hM4Di^Emx1^-dependent inhibition in the neonatal hippocampus by performing somatic current-clamp recordings. At P2–5, we found that bath-application of C21 (1 µM) did not significantly affect the resting membrane potential (RMP) of CA1 PCs in either *hM4Di^Emx1^* or WT mice (Fig. 2A-B) (GLMM; for detailed statistics, see Extended Data Table 1-1). We noticed a minor temporal drift in membrane resistance (R_m_) during these prolonged recordings. However, this time-dependent change was independent of genotype (interaction: P = 0.30; Fig. 2A and 2C), indicating that hM4Di^Emx1^ activation left R_m_ unaffected. We further explored potential effects of hM4Di^Emx1^ activation on intrinsic excitability during the second postnatal week (Fig. 2D), when active and passive membrane properties are considerably more mature but burst-like network activity is still prominent *in vivo* (Graf et al., 2022). We found that, at P8–12, both RMP and R_m_ were unaltered by bath-application of C21 (Fig. 2E-F). In summary, activation of hM4Di^Emx1^ did not induce detectable hyperpolarization or shunting in CA1 PCs before eye opening.

**Figure 2.**
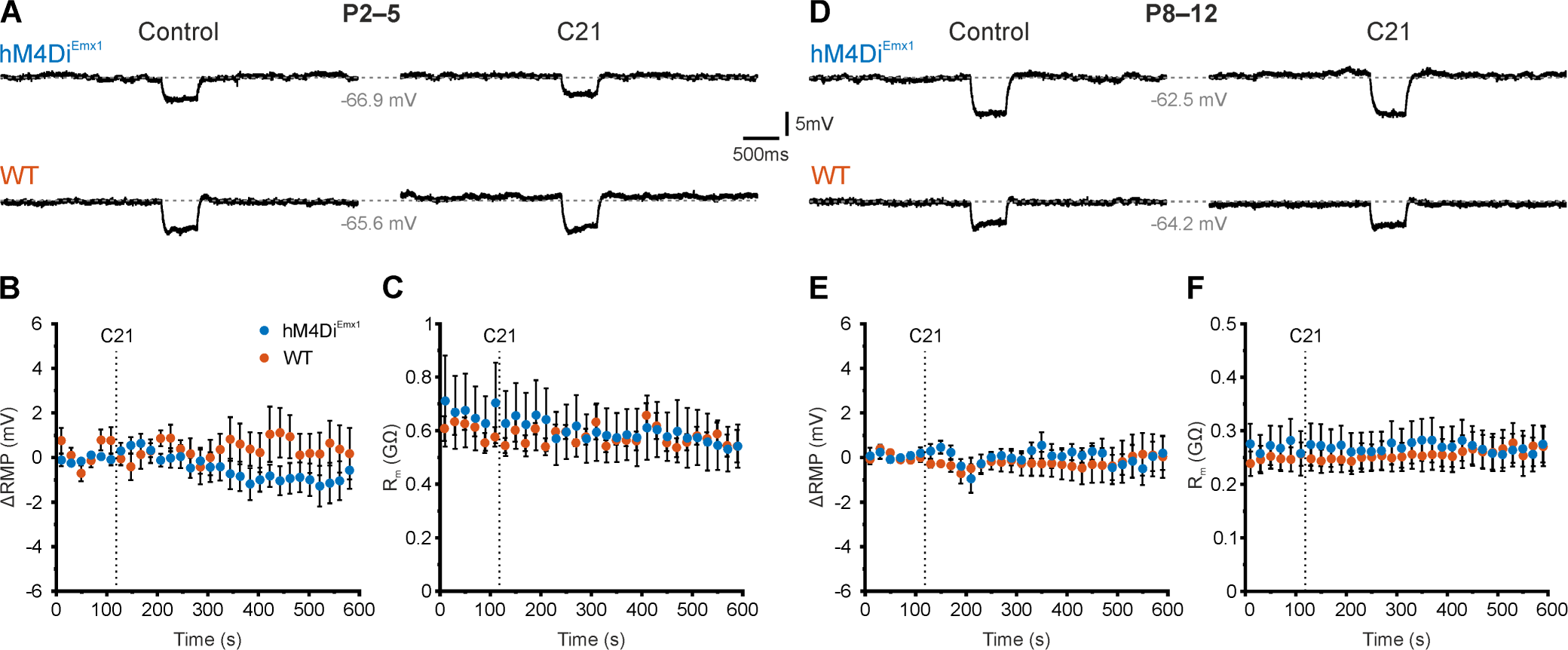
Membrane potential and membrane resistance are unaffected by *hM4Di^Emx1^* activation. ***A-C,*** Data obtained at P2–5. ***A,*** Sample current-clamp recordings (*I* = 0) of membrane potential before (*left*) and after (*right*) wash-in of C21 (1 µM). Brief test pulses (−10 pA) were used to estimate membrane resistance. ***B-C,*** Time-course of the change in resting membrane potential (ΔRMP, ***B***) and membrane resistance (R_m_, ***C***) before and during application of C21 (1 µM). The dotted lines represent application onset. ***D–F,*** The same as in ***A-C***, but at P8–12 (test pulses: −20 pA). ***B-C*** and ***E-F,*** Population data (closed symbols) are presented as mean ± SEM. For statistics, see Extended Data Table 1-1.

We next sought to extend these findings by examining the input-output relationship and characteristics of action-potential (AP) firing in CA1 PCs through current-clamp recordings in slices from *hM4Di^Emx1^* and WT mice at P2–5 (Fig. 3A-C). In each recorded PC, we applied three series of depolarizing step-current injections in the absence and presence of C21 (1 µM). Immediately prior to each experiment, the RMP was biased to −70 mV by somatic current injection so as to exclude any contribution of differences in RMP on the analyzed properties. We found that the maximum instantaneous firing rates of CA1 PCs were unaffected by C21 in slices from both *hM4Di^Emx1^* and WT mice (Fig. 3D; for detailed statistics, see Extended Data Table 1-1). During these experiments, maximum mean firing rates (per 500-ms-long current step) showed a minor decline over time, which, however, did not significantly differ between *hM4Di^Emx1^* and WT mice (interaction: P = 0.052; Fig. 3E). In either genotype, C21 lacked significant effects on AP thresholds (Fig. 3F) or rheobase currents (Fig. 3G). We then repeated these experiments during the second postnatal week (Fig. 3H-J). As compared to neurons of the first postnatal week, CA1 PCs recorded at P8–12 exhibited substantially higher maximum instantaneous (Fig. 3K) and mean (Fig. 3L) firing rates, more negative AP thresholds (Fig. 3M) and higher rheobase currents (Fig. 3N), implying that membrane properties had undergone a considerable maturation within the period investigated here. In line with our data obtained at P2–5, however, GLMM indicated that none of the analyzed quantities was significantly affected by C21 (Fig. 3K-N and Extended Data Table 1-1).

**Figure 3.**
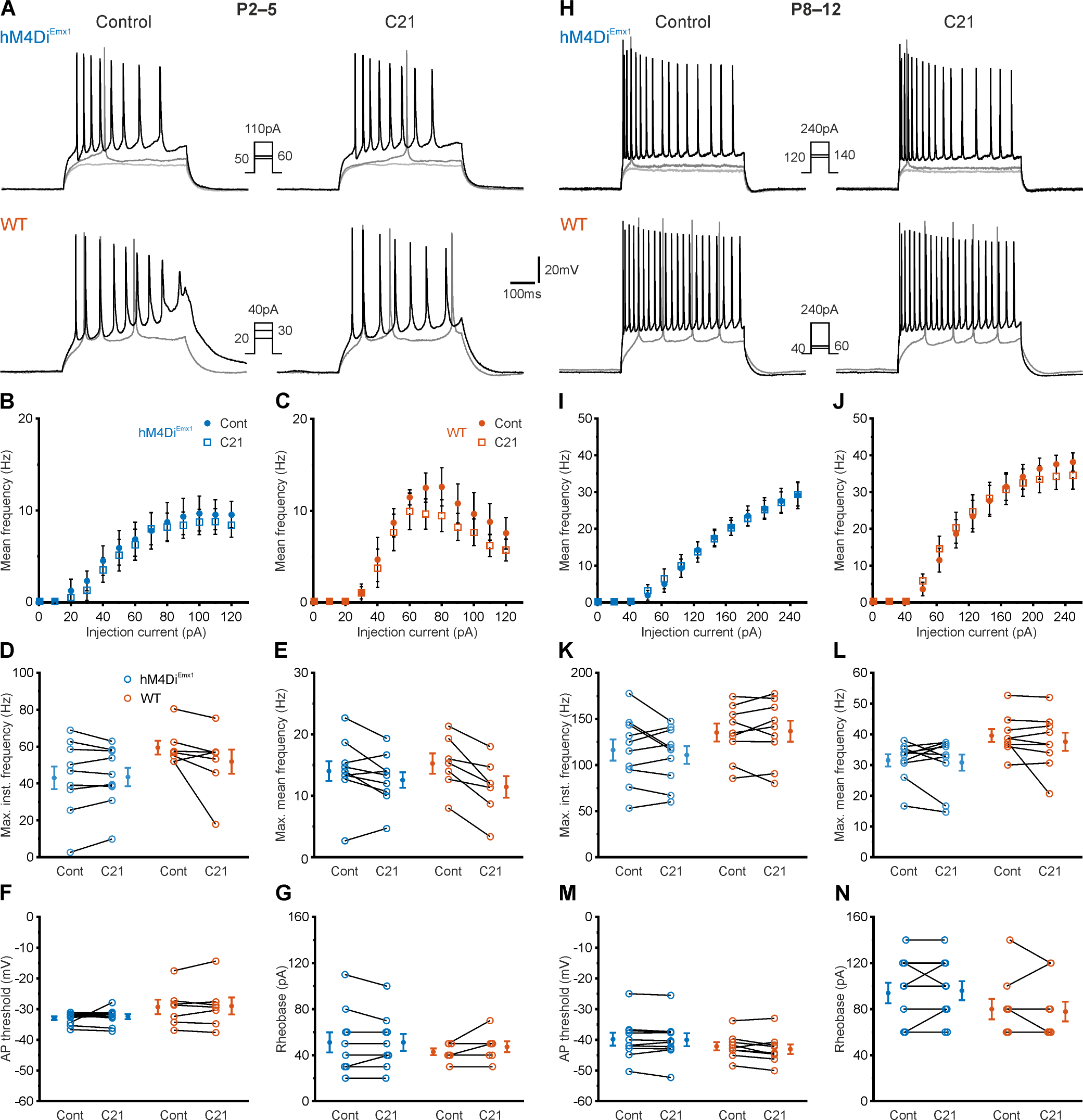
Activation of *hM4Di^Emx1^* does not affect somatic intrinsic excitability. ***A-G,*** Data obtained at P2–5. ***A,*** Sample current-clamp recordings in response to depolarizing current steps (500 ms) before (*left*) and after (*right*) wash-in of C21 (1 µM). Insets show step current amplitudes for the displayed traces (black traces correspond to maximum mean firing frequency in control, medium gray traces to rheobase current and light gray traces to the highest sub-threshold current). Recordings were performed in the continuous presence of DNQX (10 µM), APV (50 µM) and gabazine (10 µM). ***B-C,*** Input-output relationship for all cells recorded from *hM4Di^Emx1^* (***B***) and WT (***C***) mice, respectively. Population data are presented as mean ± SEM. ***D–G,*** Maximum instantaneous firing frequency (***D***), maximum mean firing frequency (***E***), AP threshold (***F***) and rheobase current (***G***). ***H-N,*** The same as in ***A-G***, but at P8–12. ***D-G*** and ***K-N***, Each open symbol represents a single cell. Population data (closed symbols) are presented as mean ± SEM. For statistics, see Extended Data Table 1-1.

Taken together, our data indicate that the observed hM4Di^Emx1^-mediated silencing of network activity in the neonatal hippocampus (Fig. 1) is unlikely to result from changes in intrinsic excitability of CA1 PCs and, thus, point to an alternative mechanism.

### hM4Di^Emx1^ activation potently suppresses synaptic glutamate release by the first postnatal week

Presynaptic G_i/o_ signaling is functional already at early developmental stages (Fukuda et al., 1993; Safiulina et al., 2005; Kirmse et al., 2008). Moreover, suppression of vesicular release was previously identified as a major mechanism of G_i/o_-based chemogenetic actuators in the adult brain (Stachniak et al., 2014; Vardy et al., 2015). We therefore investigated whether inhibition of synaptic transmission underlies the hM4Di^Emx1^-mediated silencing at the network level. To this end, we first recorded miniature excitatory postsynaptic currents (mEPSCs) in the presence of blockers of voltage-gated Na^+^ channels (tetrodotoxin, 0.5 µM) and GABA_A_ receptors (gabazine, 10 µM) (Fig. 4A). Of note, most neurons providing glutamatergic input to CA1 are derived from the *Emx1* lineage (Sando et al., 2017; Graf et al., 2021; Leprince et al., 2023). We found that, in CA1 PCs recorded from *hM4Di^Emx1^* mice at P2–5, C21 (1 µM) reduced the frequency of mEPSCs from 0.13 ± 0.03 min^−1^ to 0.05 ± 0.01 min^−1^ (Fig. 4B). Conversely, mEPSC frequency was unaffected in WT neurons, thereby excluding the possibility that hM4Di-independent actions of C21 could underlie the effect (interaction: P = 3.5 × 10^−4^; *hM4Di^Emx1^*: P = 1.1 × 10^−3^, WT: P = 0.14, simple contrasts; Fig. 4B and Extended Data Table 1-1). In addition, median mEPSC amplitudes, a measure of quantal size, remained unaltered after wash-in of C21 in either genotype (Fig. 4C), pointing to a presynaptic site of action of hM4Di^Emx1^. Again, we extended these investigations to the second postnatal week (Fig. 4D). At P8–12, akin to the above observations, C21 reduced mEPSC frequencies by about 50% selectively in *hM4Di^Emx1^* mice (Fig. 4E), without decreasing median mEPSC amplitudes (interaction: P = 0.056; Fig. 4F). Collectively, our data suggest that activation of hM4Di^Emx1^ leads to inhibition of vesicle release at glutamatergic synapses impinging on CA1 PCs, which in turn can explain the observed suppression of synchronized network activity.

**Figure 4.**
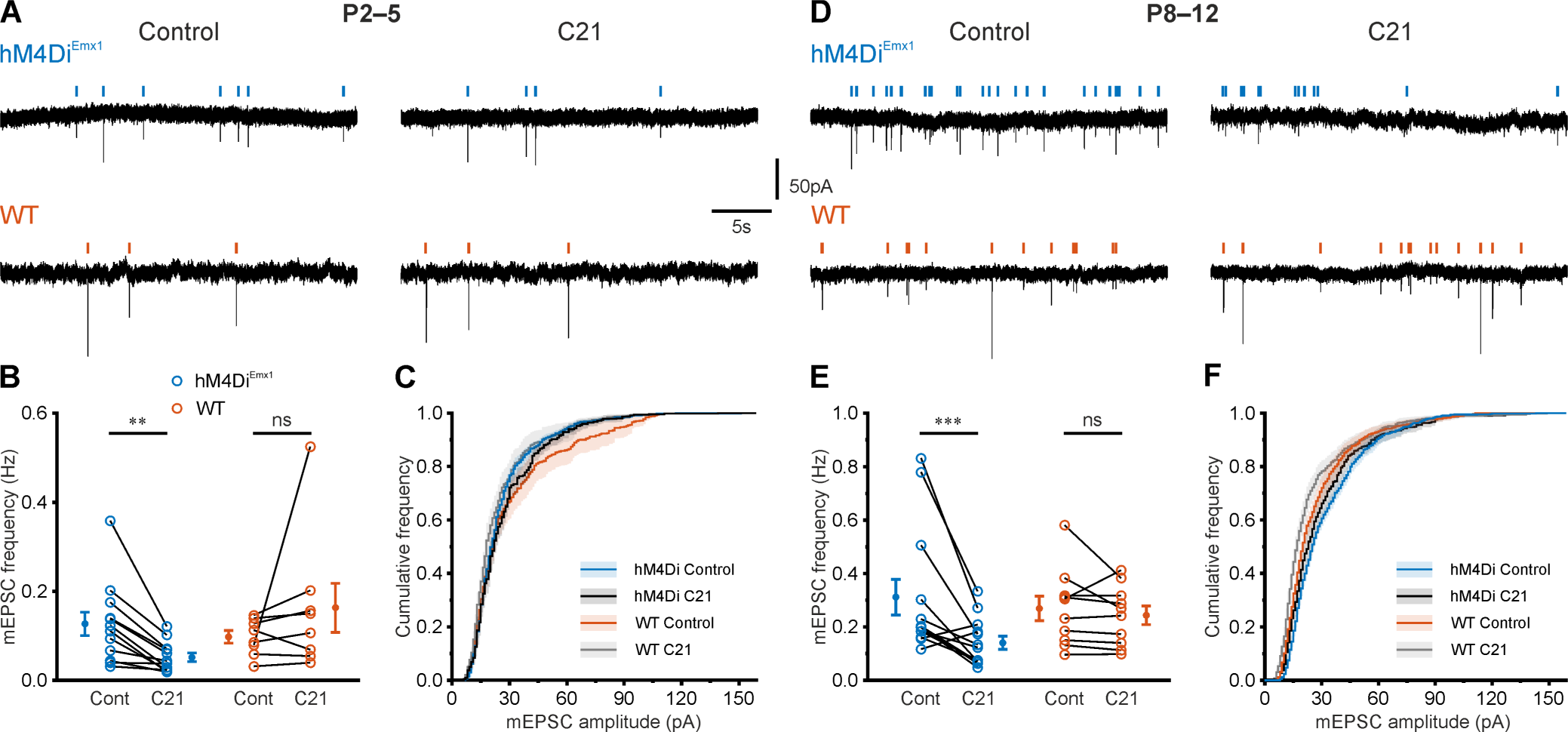
Activation of *hM4Di^Emx1^* inhibits mEPSCs in CA1 PCs. ***A-C,*** Data obtained at P2–5. ***A,*** Sample traces of mEPSCs in the absence (*left*) or presence (*right*) of C21 (1 µM). Recordings were performed in the continuous presence of TTX (0.5 µM) and gabazine (10 µM). Vertical ticks indicate detected events. ***B,*** C21 reduced mean mEPSC frequencies in PCs from *hM4Di^Emx1^*, but not WT, mice. ***C,*** mEPSC amplitude distributions. Mean (solid lines) ± SEM (shaded areas). ***D-F,*** The same as in ***A-C***, but at P8–12. ***B*** and ***E,*** Each open symbol represents a single cell. Population data (closed symbols) are presented as mean ± SEM. *** P < 0.001, ** P < 0.01, ns – not significant (simple contrasts). For statistics, see Extended Data Table 1-1.

To corroborate this conclusion, we next examined to what extent G_i/o_ signaling controls glutamate release at Schaffer collateral synapses, which provide a major excitatory drive to CA1. To this end, EPSCs evoked by electrical stimulation of afferents in *stratum radiatum* (eEPSCs) were recorded from CA1 PCs of *hM4Di^Emx1^* mice in the presence of gabazine (10 µM) (Fig. 5A). At P2–5, bath-application of C21 (1 µM) was found to massively decrease mean eEPSC amplitudes from 34.1 ± 8.5 pA to 1.7 ± 1.3 pA (Fig. 5B). This was associated with an increase in eEPSC failure rate from 25 ± 6% to 92 ± 4% (Fig. 5C), supporting the conclusion that the effect is presynaptic in origin. In these experiments, paired-pulse stimulation frequently induced long-latency polysynaptic bursts of EPSCs, reflecting network disinhibition due to the blockade of GABA_A_ receptors (arrows in Fig. 5A). We therefore considered the possibility that these EPSC bursts *per se* could change synaptic strength, i.e. independently of C21. To address this point, we performed an additional set of experiments, in which vehicle was applied instead of C21 (Fig. 5A). We found that both mean amplitudes (Fig. 5B) and failure rates (Fig. 5C) of eEPSCs were stable over time. For each parameter analyzed, GLMM revealed a significant interaction between treatment and dataset (eEPSC amplitude: P = 5.3 × 10^−9^; eEPSC failure rate: P = 2.9 × 10^−7^), and *post-hoc* tests confirmed that the suppression of eEPSCs was specific to C21 (Fig. 5B-C; for detailed statistics, see Extended Data Table 1-1). Likewise, at P8–12, activation of hM4Di^Emx1^ by bath-application of C21 profoundly reduced mean eEPSC amplitudes and increased eEPSC failure rates, both of which were not observed when vehicle was applied instead of C21 (Fig. 5D-F). Taken together, these data demonstrate that activation of G_i/o_ signaling through hM4Di^Emx1^ potently suppresses synaptic glutamate release at Schaffer collateral synapses on CA1 PCs by the first postnatal week.

**Figure 5.**
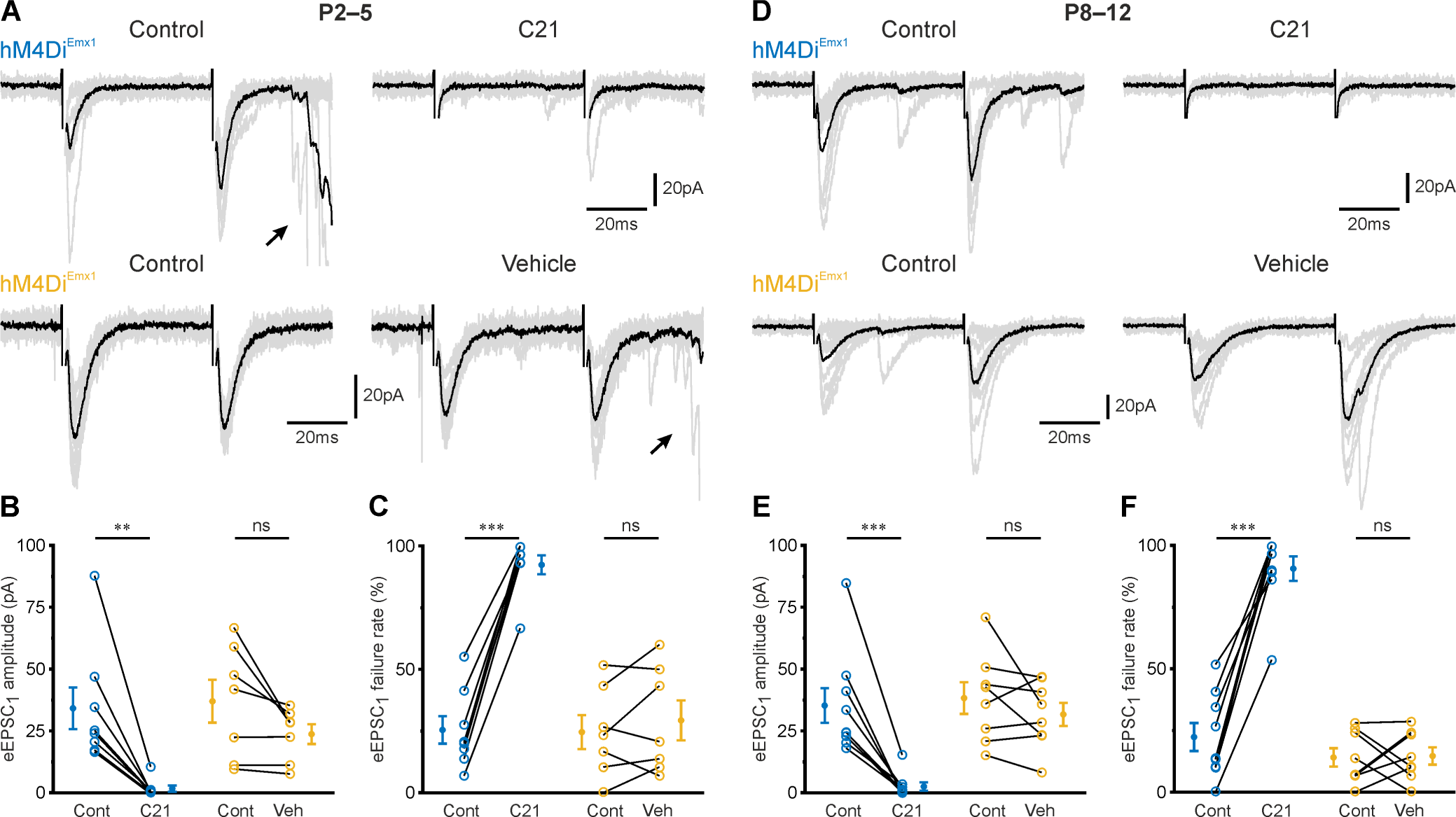
*hM4Di^Emx1^* activation suppresses evoked glutamate release at Schaffer collateral synapses. ***A-C,*** Data obtained at P2–5. ***A,*** Sample EPSCs evoked by stimulation of Schaffer collaterals in the absence (*top-left*) or presence (*top-right*) of C21 (1 µM). In a separate dataset (*bottom*), vehicle (instead of C21) was applied to examine the temporal stability of evoked responses. Recordings were performed in the continuous presence of gabazine (10 µM). Stimulus artefacts were clipped for clarity. Arrows indicate long-latency polysynaptic bursts of EPSCs. Gray traces correspond to ten successive trials, black traces represent their means. ***B-C,*** Mean amplitudes (***B***) and failure rates (***C***) of EPSCs evoked by the first pulse (eEPSC_1_). ***D-F,*** The same as in ***A-C***, but at P8–12. ***B-C*** and ***E-F,*** Each open symbol represents a single cell. Population data (closed symbols) are presented as mean ± SEM. *** P < 0.001, ** P < 0.01, ns – not significant (simple contrasts). For statistics, see Extended Data Table 1-1.

### Presynaptic G_i/o_ signaling is an actuator of immature hippocampal network dynamics in vivo

Our data obtained so far indicate that the hM4Di^Emx1^-mediated inhibition of synaptic glutamate release alone suffices to virtually abolish GDPs in acute slices and therefore point to presynaptic G_i/o_ signaling as an actuator of neuronal synchrony in the developing hippocampus. However, the *in vivo* relevance of these findings remains unclear. We therefore performed local field potential (LFP) recordings in *stratum radiatum* of CA1 in head-fixed mice. Experiments were conducted under nitrous oxide anesthesia at P3–5. At this developmental stage, CA1 PCs are incapable of sustaining persistent firing activity but instead generate intermittent population bursts reminiscent of GDPs in slices (Leinekugel et al., 2002; Dard et al., 2022; Graf et al., 2022). In accordance with previous studies, LFP activity at P3–5 was discontinuous and dominated by early sharp waves (eSPWs; Fig. 6A-B) (Valeeva et al., 2019; Gainutdinov et al., 2023). Less consistently, we also detected short-lasting LFP oscillations in the theta-to-beta frequency range in several of the analyzed mice (Fig. 6A) (Graf et al., 2021). For quantification of hM4Di^Emx1^ effects, we computed the LFP bandpower (8–40 Hz) in non-overlapping 5-min-long intervals and normalized it to the bandpower of the technical noise (see Methods for details). In *hM4Di^Emx1^* mice, a single dose of systemically applied C21 (3 mg/kg s.c.) reduced the normalized bandpower to approximately noise levels, implying an almost complete suppression of LFP activity (Fig. 6C-D). This effect set in within minutes, confirming the favorable bioavailability of C21 when administered through subcutaneous injection in neonatal mice (Thompson et al., 2018; Jendryka et al., 2019; Kostka and Hanganu-Opatz, 2023). The suppression of LFP activity lasted for at least two hours (Fig. 6C) and was strictly dependent on activation of hM4Di^Emx1^, since no decrease in normalized bandpower was observed upon application of either vehicle in *hM4Di^Emx1^* pups or C21 in WT mice at 30–60 min *post* injection (interaction: P = 2.5 × 10^−9^). Likewise, eSPWs were virtually abolished after injection of C21 in *hM4Di^Emx1^* mice (control: 2.1 ± 0.3 min^−1^, C21: 0.0 ± 0.0 min^−1^; Fig. 6B and 6E; for detailed statistics, see Extended Data Table 1-1). While the normalized bandpower in WT mice remained unaffected, C21 reduced eSPW rates, although to a lesser extent than in *hM4Di^Emx1^* pups.

**Figure 6.**
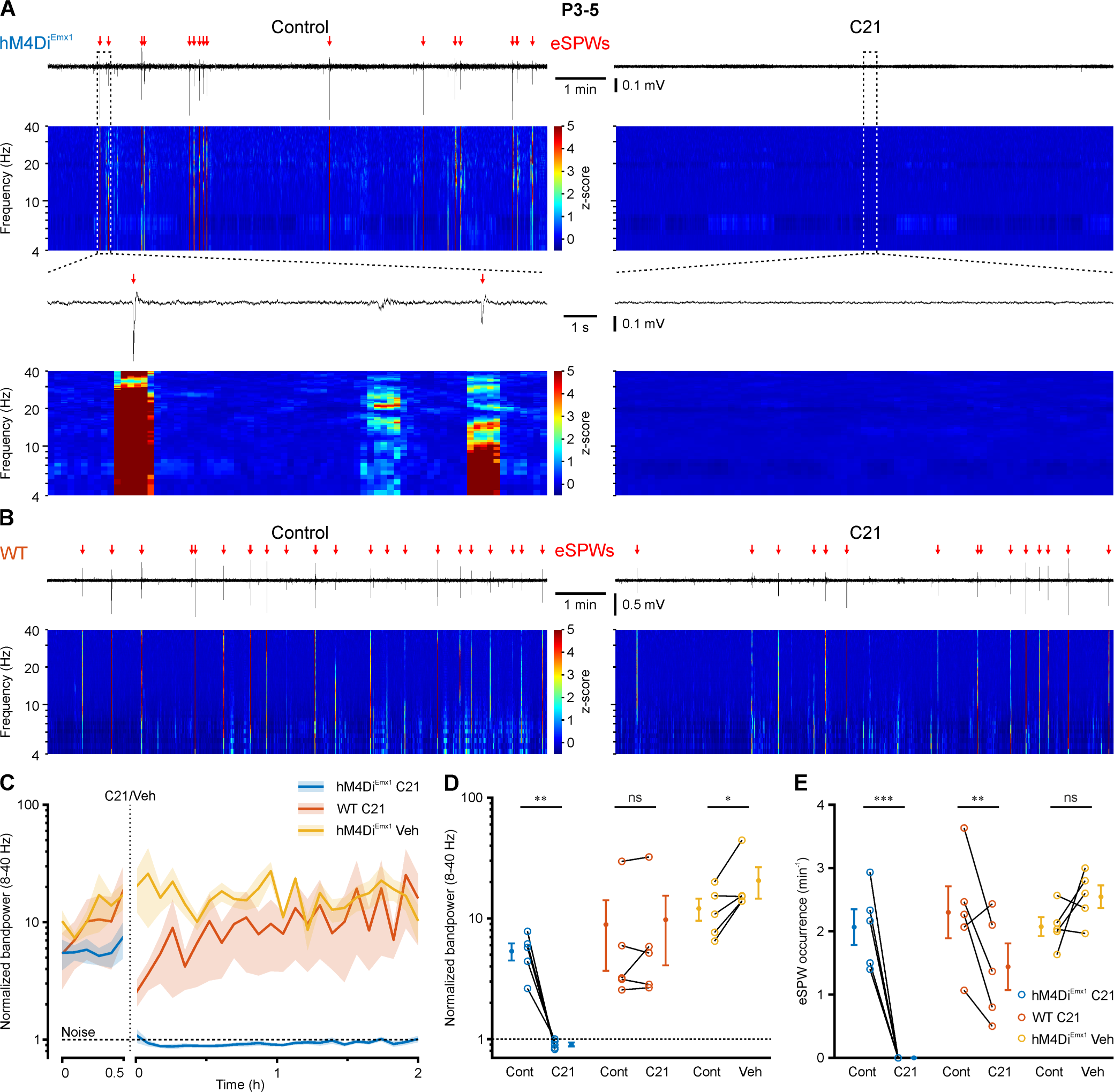
Activation of *hM4Di^Emx1^* effectively inhibits early sharp waves (eSPWs) in CA1 *in vivo* at P3–5. ***A,*** Sample LFP recording and time-aligned spectrogram (4–40 Hz, window length: 1 s, overlap 80%, color corresponds to z-score across frequency domain) from the dorsal CA1 of an *hM4Di^Emx1^* mouse before (*left*) and after (*right*) injection of C21 (3 mg/kg s.c.). Red arrows indicate eSPWs. *Bottom*, boxed regions at higher temporal resolution. ***B,*** The same as in ***A*** (*top*), but for a WT animal. ***C,*** Time-course of bandpower (8–40 Hz) normalized to the technical noise (see Methods). Mean (solid lines) ± SEM (shaded areas). Note the immediate effect of C21 in *hM4Di^Emx1^* animals, reducing the bandpower to the noise level (dashed line). ***C-D***, Y axis is shown in log scale. ***D-E,*** Normalized bandpower (8–40 Hz) (**D**) and eSPW occurrence frequency (**E**) for *hM4Di^Emx1^* and WT animals before and after injection of C21 or vehicle. Each open symbol represents a single animal. Population data (closed symbols) are presented as mean ± SEM. *** P < 0.001, ** P < 0.01, * P < 0.05, ns – not significant (simple contrasts). For statistics, see Extended Data Table 1-1.

We finally asked whether hM4Di^Emx1^-induced presynaptic inhibition of synaptic glutamate release can effectively silence network activity at P10–12. By this time, discrete eSPW events had been largely replaced by more complex, mostly discontinuous, oscillatory LFP activity patterns in the theta-beta range (Fig. 7A), reflecting a profound developmental increase in synaptic inputs to CA1. In *hM4Di^Emx1^* mice, subcutaneous injection of C21 reduced the normalized bandpower by approximately one order of magnitude, with little recovery during the subsequent two-hour recording period (Fig. 7A and 7C). Maximum suppression was already present 15 min after injection. Conversely, no apparent changes in normalized bandpower were evident upon application of either vehicle in *hM4Di^Emx1^* pups or C21 in WT mice (Fig. 7B-C). Accordingly, GLMM revealed a significant interaction term (dataset × treatment: P = 2.1 × 10^−12^), and *post-hoc* testing confirmed that the suppression of neuronal activity by C21 was restricted to *hM4Di^Emx1^* mice (Fig. 7C; for detailed statistics, see Extended Data Table 1-1). Taken together, our data demonstrate that hM4Di^Emx1^-induced synaptic suppression of glutamate release can impose powerful inhibition on synchronized network activity in the developing hippocampus before eye opening.

**Figure 7.**
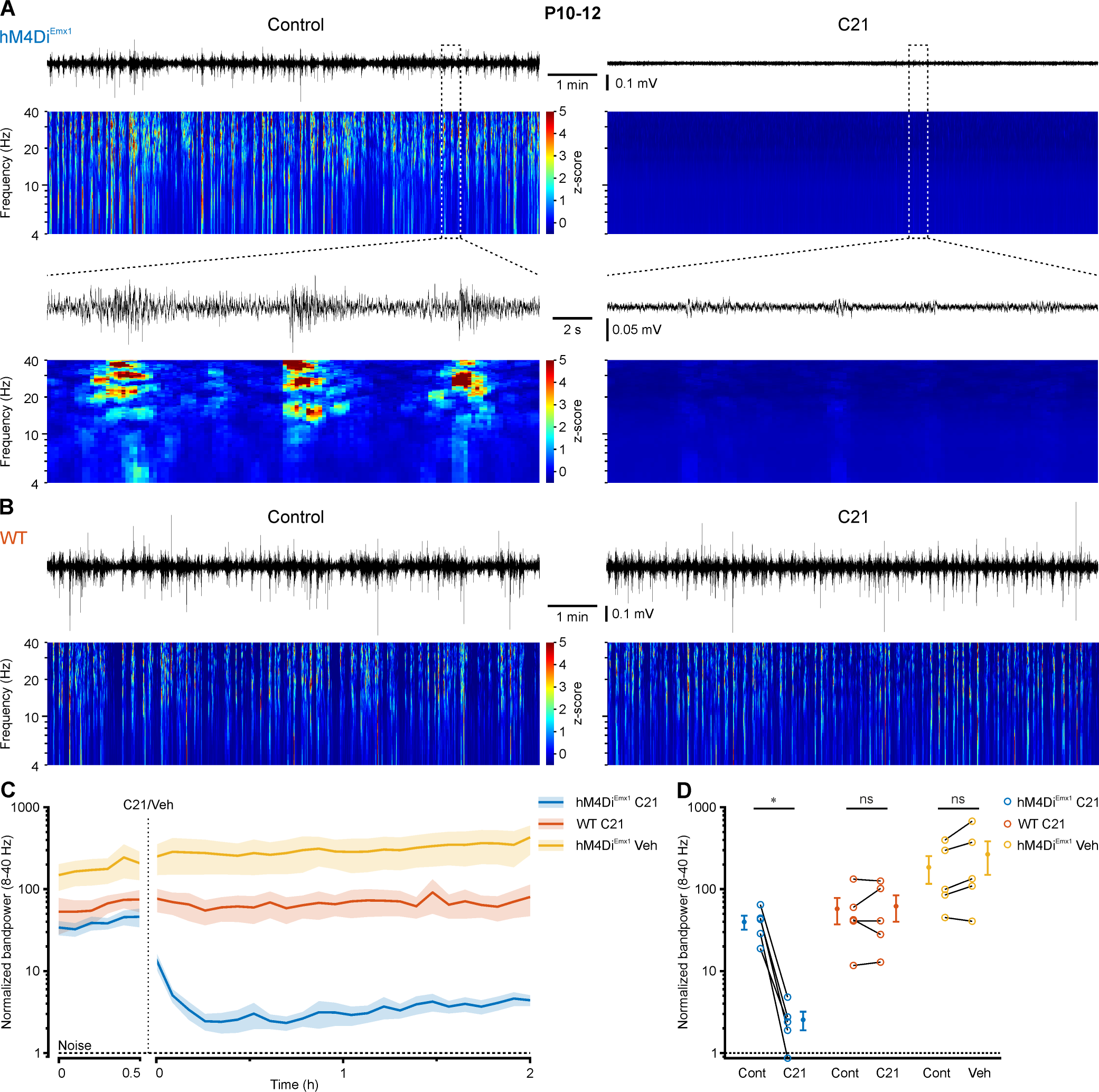
Effective silencing of hippocampal network oscillations through *hM4Di^Emx1^* activation in CA1 *in vivo* at P10–12. ***A,*** Sample LFP recording and time-aligned spectrogram (4–40 Hz, window length: 1 s, overlap 80%, color corresponds to z-score across frequency domain) from the dorsal CA1 of a *hM4Di^Emx1^* mouse before (*left*) and after (*right*) wash-in of C21 (3 mg/kg s.c.). *Bottom*, boxed regions at higher temporal resolution. ***B,*** Same as in **A** (*top*), but for a WT animal. ***C,*** Time course of bandpower (8–40 Hz) normalized to the technical noise (see Methods) for control (30 min) and after C21 injection. Y axis is shown in log scale. Mean (solid lines) ± SEM (shaded areas). Note that the effect peaked about 15 min after C21 injection in *hM4Di^Emx1^* animals and recovered slowly. The red dashed line indicates the electrode noise level. ***D,*** Normalized bandpower (8–40 Hz) for *hM4Di^Emx1^* and WT animals before and after injection of C21 or vehicle. Each open symbol represents a single animal. Population data (closed symbols) are presented as mean ± SEM. * P < 0.05, ns – not significant (simple contrasts). For statistics, see Extended Data Table 1-1.

## Discussion

### Generative mechanisms of network bursts in the developing CA1

Network bursts in the developing hippocampus are generated in a relatively all-or-none manner (Prida and Sanchez-Andres, 1999; Gainutdinov et al., 2023). Theoretical work suggests that this behavior arises from a transient, unstable network state evoked by supra-threshold excitatory input, leading to a profound amplification of neuronal firing rates (Rahmati et al., 2017). To further characterize the generative mechanisms underlying synchronized activity, we here employed a transgenic approach (*hM4Di^Emx1^* mice) (Gorski et al., 2002; Zhu et al., 2016) to target the entire population of *Emx1*-lineage cells from embryonic stages onwards. We examined the effect of hM4Di^Emx1^ activation *in vitro* and found that it abolishes GDPs at P2–5 (Fig. 1). Likewise, a single systemic injection of the DREADD agonist C21 completely blocked eSPWs as observed in *in vivo* LFP recordings from neonatal *hM4Di^Emx1^* mice (Fig. 6). Thus, glutamatergic neurons of the *Emx1*-lineage contribute most to the transient instability underlying the supra-threshold amplification of firing rates during network bursts in CA1. This further implies that neither GDPs *in vitro* nor eSPWs *in vivo* are sustained by depolarizing GABAergic (*Emx1^−^*) neurons alone, despite their partially excitatory action (Flossmann et al., 2019; Murata and Colonnese, 2020; Graf et al., 2021). In addition, our data are fully compatible with the previous conclusion that somatosensory feedback, in the form of eSPWs, is relayed to CA1 via *Emx1^+^* glutamatergic projections from the entorhinal cortex (Valeeva et al., 2019; Leprince et al., 2023), whereas glutamatergic input from *Emx1^−^* sources, such as from the ventral midline thalamus (Leprince et al., 2023), modulates rather than sustains synchronized activity in developing CA1. Similar considerations apply to the second postnatal week, when short-lasting events (i.e., eSPWs) had been largely replaced by discontinuous oscillatory LFP activity (Fig. 7), consistent with the single-cellular dynamics of network activity in developing CA1 *in vivo* (Graf et al., 2022). In methodological terms, our data demonstrate that transgenic expression of hM4Di enables effective suppression of spontaneous synchronized activity already shortly after birth, positioning it as a valuable and non-invasive alternative to viral transfection or *in utero* electroporation for developmental studies (Zhu et al., 2016).

### Cellular mechanism and efficacy of hM4Di^Emx1^ in neonatal CA1 neurons

G_i_-DREADDs including hM4Di are normally localized to both neuronal cell bodies and axons (Mahler et al., 2014). Accordingly, in the adult brain, G_i_-DREADD-based neuronal silencing results from the combined effects of (i) hyperpolarization and increased membrane conductance in somatodendritic domains and the (ii) suppression of vesicle release from presynaptic terminals (Armbruster et al., 2007; Stachniak et al., 2014; Vardy et al., 2015). We here examined the efficacy of hM4Di^Emx1^-mediated silencing in immature hippocampal neurons during the neonatal period, when GIRK channel expression is low (Chen et al., 1997; Ehrengruber et al., 1997) and endogenous G_i/o_-protein-coupled receptors are ineffective in mediating hyperpolarization in CA1 neurons (Nurse and Lacaille, 1999; Verheugen et al., 1999). Unlike what is observed in adult neurons, we discovered that activation of hM4Di^Emx1^ did not induce hyperpolarization or shunting (Fig. 2), thereby leaving evoked AP firing in somatic current-clamp recordings unaffected (Fig. 3). Conversely, activation of hM4Di^Emx1^ effectively suppressed glutamatergic neurotransmission, resembling previous observations in adult neurons. This synaptic silencing is likely of presynaptic origin, since hM4Di^Emx1^ caused a selective reduction in the frequency, but not amplitude, of mEPSCs (Fig. 4) as well as a substantial increase in the failure rate of eEPSCs (Fig. 5). The observed dichotomy supposedly reflects differences in the signaling cascades engaged by G_i/o_-βγ dimers, which, in presynaptic terminals, inhibit evoked and spontaneous (miniature) release chiefly by interacting with SNARE proteins and P/Q- and N-type Ca^2+^ channels (Herlitze et al., 1996; Zurawski et al., 2019; Alten et al., 2022), as opposed to activating GIRK channels in somata and dendrites. We found that activating G_i/o_ signaling via hM4Di^Emx1^ caused an almost complete suppression of glutamatergic synaptic responses evoked in *stratum radiatum* (i.e., putative Schaffer collaterals) (Fig. 5B), indicating that presynaptic silencing is highly efficacious by P2–5. The exclusively presynaptic effect is in line with previous studies showing that G_i/o_ signaling matures earlier at pre-compared to postsynaptic sites to inhibit neurotransmitter release by the time of birth (Gaiarsa et al., 1995; Caillard et al., 1998; Nurse and Lacaille, 1999). It is important to consider that over timescales longer than those addressed in this study, G_i/o_-dependent actuators are anticipated to exert additional influence on canonical cAMP-protein kinase A pathways. These pathways have previously been implicated in, for instance, the synaptic incorporation of AMPA receptors (Esteban et al., 2003), postsynaptic long-term potentiation (Yasuda et al., 2003), presynaptic long-term plasticity (Sakaba and Neher, 2003; Atwood et al., 2014; Shahoha et al., 2022), synapse formation (Liang et al., 2021) as well as neuronal migration and neurite growth (Bony et al., 2013). Taken together, we demonstrate that hM4Di can serve as a valuable presynaptic silencer in mice shortly after birth. Our findings hold potential significance for experiments utilizing hM4Di, other G_i_-DREADDs such as KORD (Vardy et al., 2015) or likewise G_i/o_-dependent optogenetic actuators (Mahn et al., 2021) for neuronal inhibition at early developmental stages.

### Presynaptic G_i/o_ signaling as an actuator of developing hippocampal network dynamics

Our analyses lead us to hypothesize that, prior to eye opening, hippocampal network activity is under tight control of G_i/o_ signaling acting on presynaptic terminals of glutamatergic excitatory neurons. In the theoretical framework mentioned above, a reduction in quantal content of evoked glutamate release raises the network’s amplification threshold, which effectively renders network burst generation less likely (Flossmann et al., 2019). During the neonatal period, GABA_B_R-mediated postsynaptic signaling is not yet functional (Nurse and Lacaille, 1999; Verheugen et al., 1999), and the chloride extrusion capacity of immature neurons is low (Spoljaric et al., 2017). Therefore, G_i/o_-dependent presynaptic silencing could act as a powerful inhibitory constraint on CA1 activity – at developmental stages when postsynaptic GABAergic inhibition is weak (for review see Kirmse and Zhang, 2022). Currently, a major open question relates to the identity and dynamics of endogenous transmitters mediating presynaptic silencing through G_i/o_ pathways. Earlier studies identified GABA and adenosine (acting via GABA_B_ and adenosine A1 receptors, respectively) as candidates released by local neurons and/or glial cells (Gaiarsa et al., 1995; Nurse and Lacaille, 1999; Safiulina et al., 2005). Beyond that, neuromodulatory projections from subcortical nuclei could target terminals for the presynaptic silencing of excitation. While it is increasingly understood how G_i/o_-dependent neuromodulators, such as acetylcholine, dopamine, noradrenaline, and serotonin, shape mnemonic functions through the control of synaptic signaling and plasticity in the adult CA1 (Palacios-Filardo and Mellor, 2019), their developmental emergence and functions are mostly unclear. A better understanding of G_i/o_-dependent presynaptic silencing is clinically relevant, as many therapeutic and recreational drugs modulating G_i/o_ pathways can interfere with brain development during pregnancy, thereby potentially causing long-lasting cognitive impairments in the offspring (Silva et al., 2013; Fazeli et al., 2017). Exploring whether presynaptic G_i/o_-targeting neuromodulation could serve as a brain state-dependent gating mechanism for neuronal dynamics in the developing hippocampus therefore presents an interesting avenue for research.

## Acknowledgments

This work was funded by the Deutsche Forschungsgemeinschaft (DFG, German Research Foundation – Research Grants: KI 1816/6-1 #442107075, KI 1816/7-1 #448069679 to K.K.; HO 2156/5-1 #442107075, HO 2156/6-1 #448069679 to K.H.; Research Unit 3004: KI1816/5-1 #432559020, KI 1816/9-1 #415914819 to K.K.; Major Research Instrumentation: INST 93/1103-1 FUGG #502670664 to K.K.) and the Thüringer Aufbaubank (2017 FGI 0020 to K.H.). We thank Ina Ingrisch and Maria Oppmann for excellent technical assistance.

## Author Contributions

Conceptualization, K.K., K.H., J.G.; Methodology, J.G., T.F., K.K.; Formal Analysis, A.S., J.G., K.K.; Investigation, A.S., J.G., K.K.; Writing – Original Draft, K.K., J.G., A.S., K.H.; Writing – Review & Editing, K.K., J.G., K.H., A.S., T.F.; Supervision, K.K., K.H.; Funding Acquisition, K.K., K.H.

## Extended Data

**Table 1-1.**
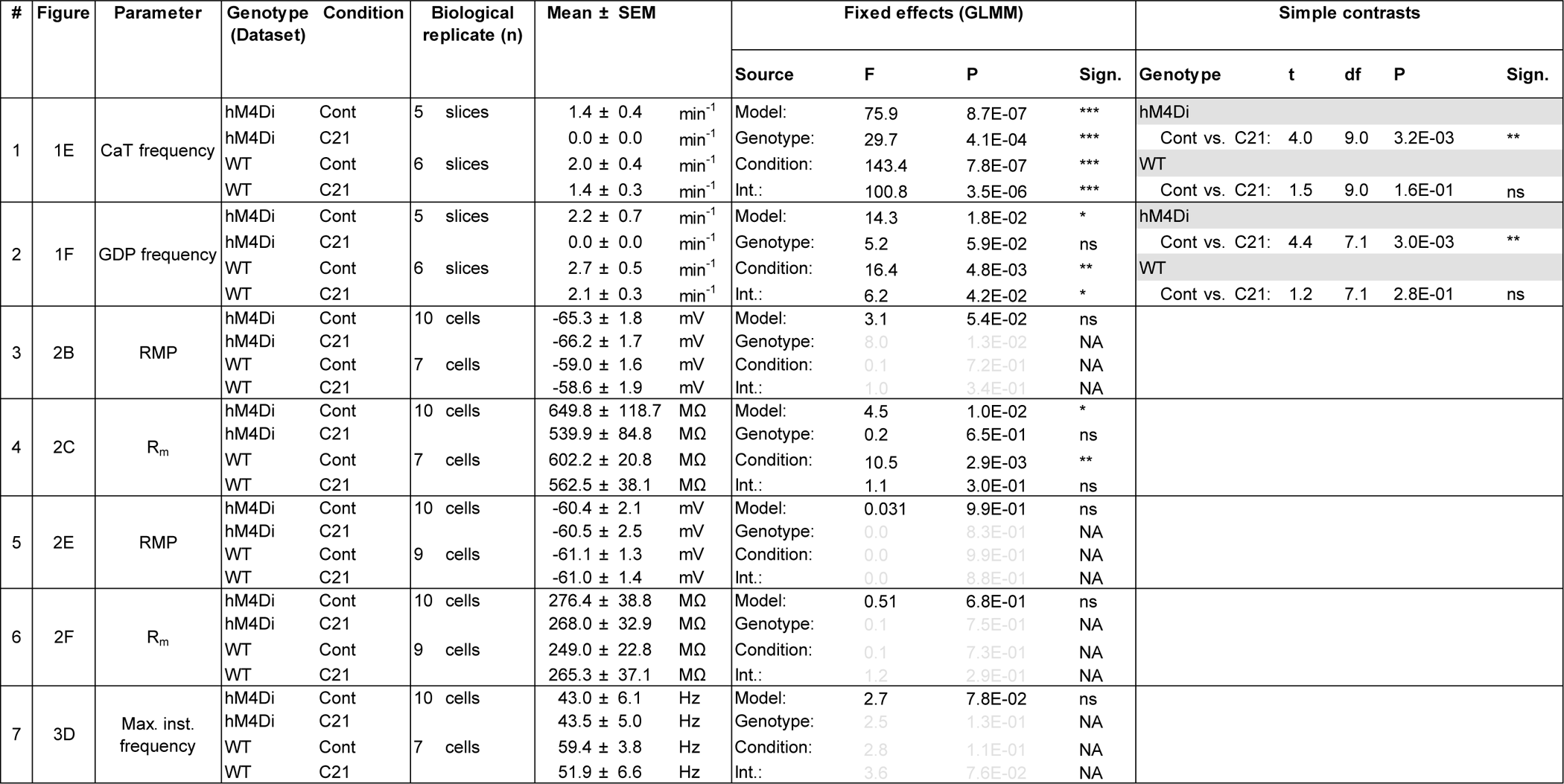
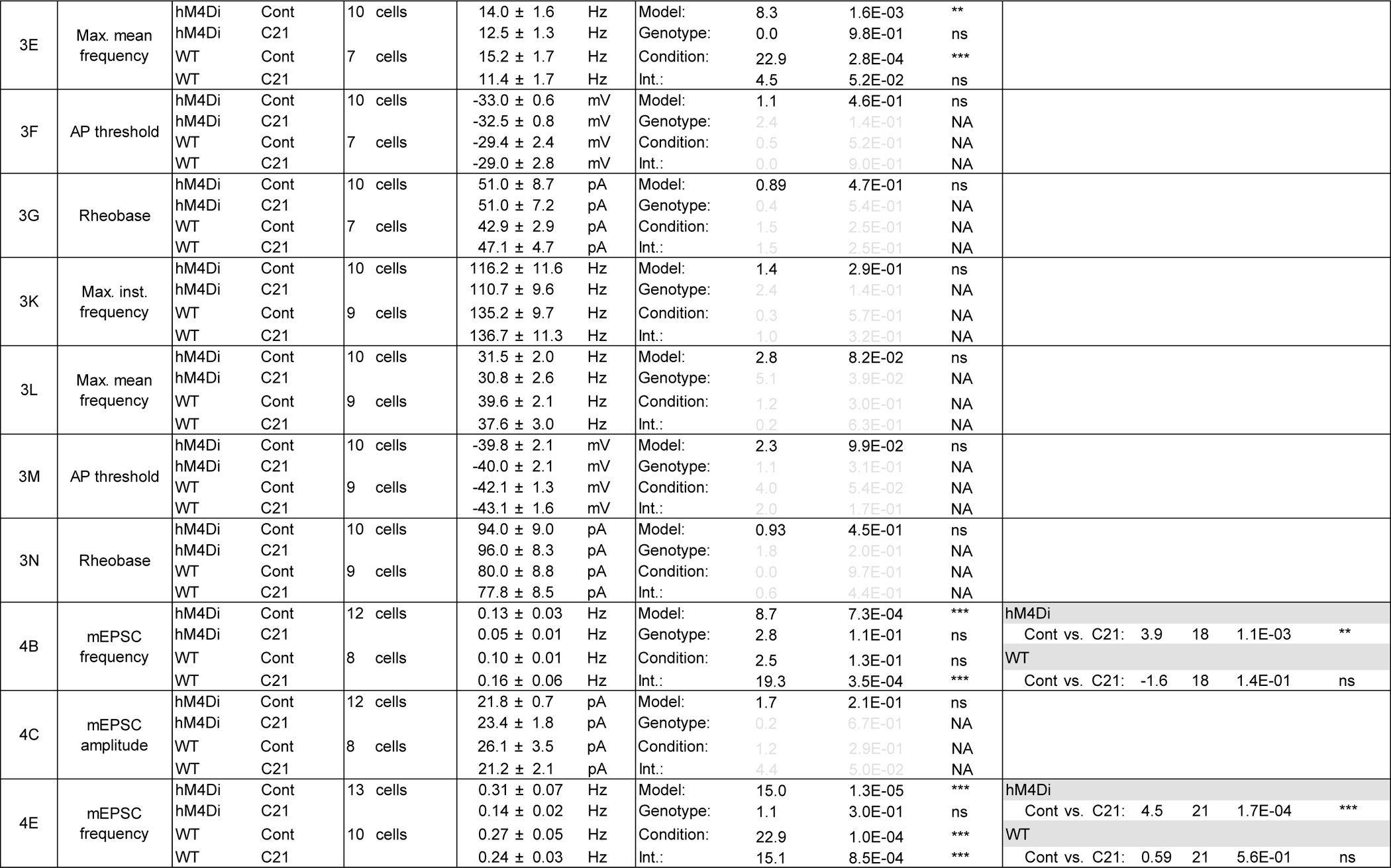
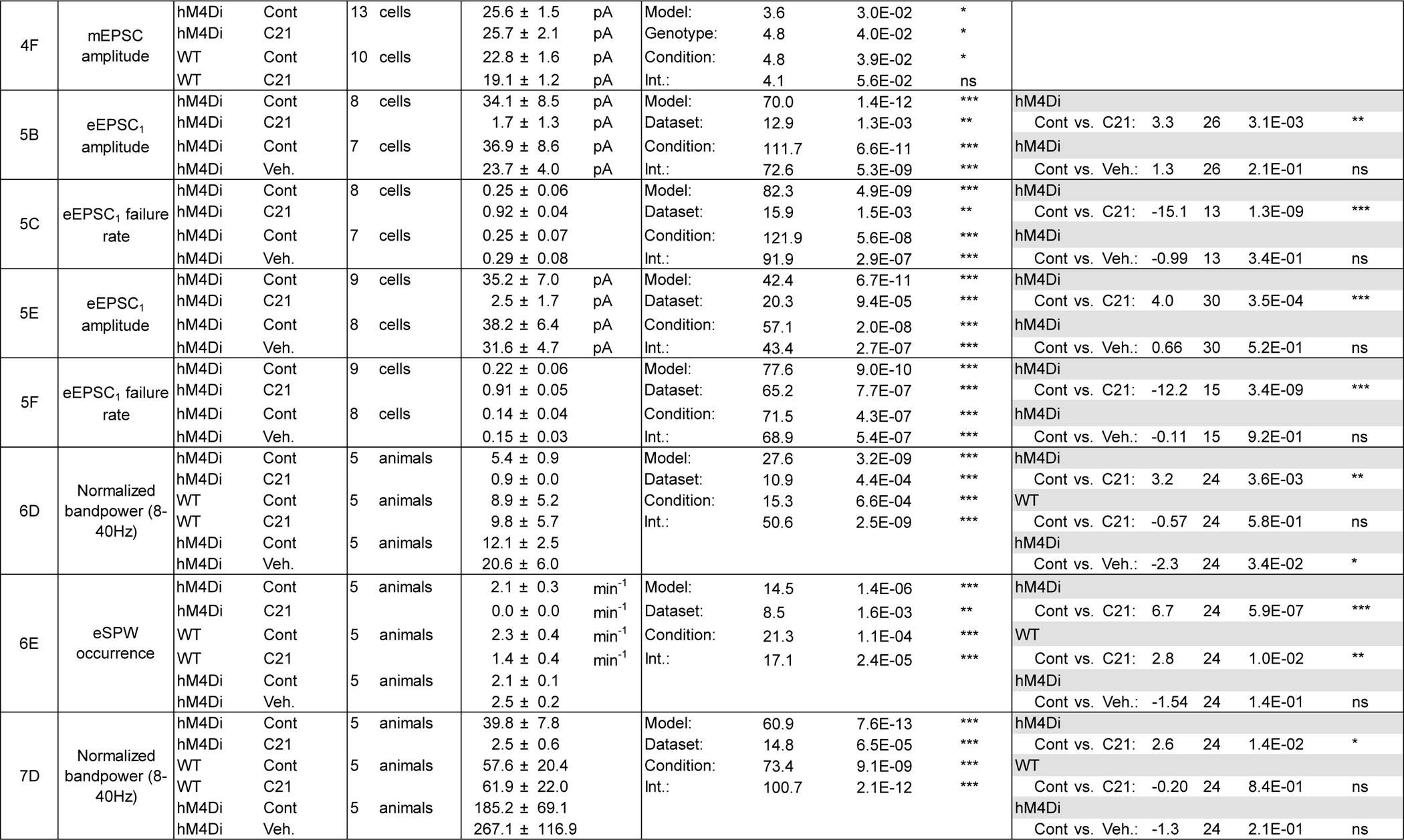
Extended Data table supporting Figures 1–7: Descriptive and inductive statistics. GLMM, generalized linear mixed model. *** P < 0.001, ** P < 0.01, * P < 0.05, ns – not significant.

## Materials and Methods

### Key Resources Table

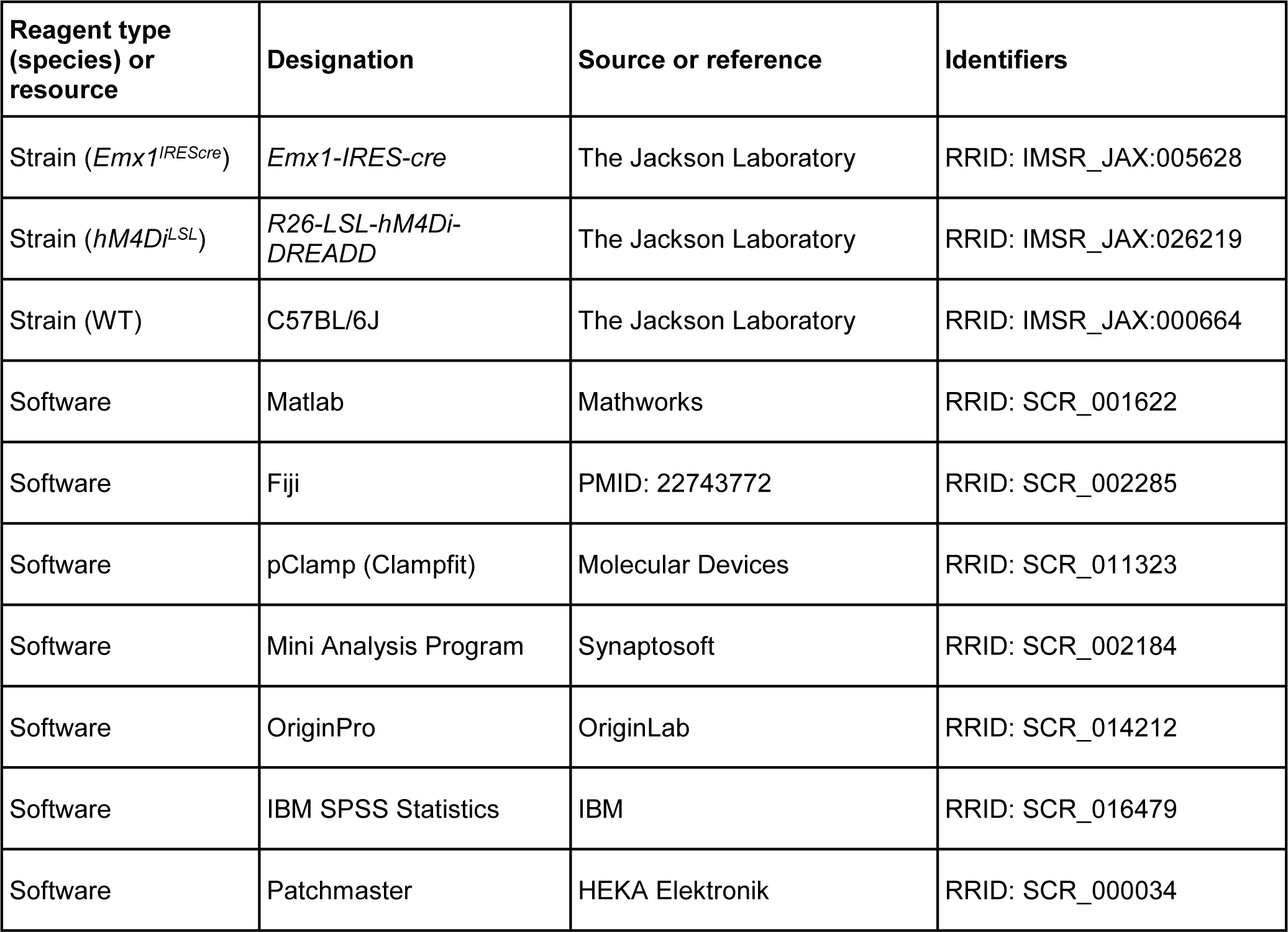

## Animals

All animal procedures were performed with approval of the local governments (Thüringer Landesamt für Verbraucherschutz, reference no.: 02-012/16; Regierung von Unterfranken, Arbeitsbereich Tierschutz) and complied with European Union norms (Directive 2010/63/EU). Animals were housed in standard cages with *ad libitum* access to food and water. *Emx1^IREScre^* (#005628) and *hM4Di^LSL^* (#026219) mice were originally obtained from The Jackson Laboratory and maintained on a C57BL/6J background. Double heterozygous offspring (referred to as *hM4Di^Emx1^ mice*) was used for experiments. C57BL/6J mice served as WT controls. Mice of either sex were used.

### Preparation of acute brain slices

Animals were decapitated under deep isoflurane anesthesia. The brain was quickly removed and transferred into ice-cold saline containing (in mM): 125 NaCl, 4 KCl, 10 glucose, 1.25 NaH_2_PO_4_, 25 NaHCO_3_, 0.5 CaCl_2_, and 2.5 or 6 MgCl_2_, gassed with carbogen (5% CO_2_, 95% O_2_; pH 7.4). Horizontal brain slices (350 µm) were cut on a vibratome and stored for at least 1 h before their use at room temperature in artificial cerebrospinal fluid (ACSF) containing (in mM): 125 NaCl, 4 KCl, 10 glucose, 1.25 NaH_2_PO_4_, 25 NaHCO_3_, 2 CaCl_2_, and 1 MgCl_2_, gassed with carbogen (pH 7.4). For recordings, slices were transferred into a submerged-type recording chamber on the microscope stage. All experiments were performed at ∼32 °C.

### Two-photon Ca^2+^ imaging in vitro

For single-cell Ca^2+^ imaging, cells were loaded with the membrane-permeable Ca^2+^ indicator Oregon Green 488 BAPTA-1 AM (OGB1) using multi-cell bolus-loading in *stratum pyramidale* of hippocampal CA1. Imaging was performed using an acousto-optic deflection (AOD) two-photon laser-scanning microscope operated by the software MES (Femto3D ATLAS, Femtonics). Fluorescence excitation at 800 nm was provided by a tunable Ti:Sapphire laser (Chameleon Ultra II, Coherent) using a 20×/1.0 NA water immersion objective (XLUMPLFLN 20XW, Olympus). Emission light was detected by photomultiplier tubes (16 bit, H11706P-40, Hamamatsu) above and below the sample. In the upper detection pathway, emission light was separated from excitation light using a primary dichroic mirror (700 nm), short-pass filtered with an IR blocker (700 nm) and further band-pass filtered (520/60 nm). In the lower detection pathway, an oil immersion condenser was used. Excitation light was blocked using an IR blocker (700 nm). The emission light was then passed through a dichroic mirror (565 nm) and band-pass filtered (520/60 nm). Signals from both photomultiplier tubes were summed up by a signal combiner and digitized. To record the activity of neurons from various z-depths within the brain slice, chessboard scanning was used, in which square scanning regions (50×50 pixels at a resolution of 1 µm/pixel) were defined at various xyz-positions. Sampling rate depended on the number of scanning regions and ranged from about 40 to 62 Hz. Spontaneous activity was recorded for 8 min before and after wash-in of C21.

### Patch-clamp recordings in vitro

Electrophysiological signals were acquired using a HEKA EPC 10 amplifier, a built-in 16-bit AD/DA board and the software Patchmaster (HEKA Elektronik). Signals were low-pass filtered at 2.9 kHz and sampled at 20 kHz. For whole-cell current-clamp recordings, glass pipettes (3– 5 MΩ) were filled with solution containing (in mM): 12 KCl, 123 K-gluconate, 5 NaCl, 0.2 EGTA, 10 HEPES, 1.8 Mg-ATP, 0.3 Na-GTP, 0.1 spermine, 10 phosphocreatine (pH 7.3). Ionotropic glutamate and GABA receptor antagonists (10 µM DNQX, 50 µM APV, 10 µM gabazine) were added to the ACSF to eliminate recurrent excitation and to minimize synaptic noise. Resting membrane potential was measured for a total of 10 min at zero current. Once per 20 s, a test pulse (duration: 500 ms; amplitude: −10 pA at P2–5, −20 pA at P8–12) was applied to estimate membrane resistance. C21 (1 µM) was applied after 2 min of control recording. AP firing characteristics were measured by applying a series of step currents (each 500 ms) of variable amplitude (P2–5: from 0 to 120 pA in increments of 10 pA; P8–12: from 0 to 240 pA in increments of 20 pA). Three series were recorded in each condition, both control and C21. Immediately prior to the first series in each condition, the membrane potential was biased to −70 mV. For recording mEPSCs, whole-cell voltage-clamp measurements were performed at a holding potential of −70 mV. Recording pipettes were filled with solution containing (in mM): 145 KCl, 5 NaCl, 0.2 EGTA, 10 HEPES, 1.8 Mg-ATP, 0.3 Na-GTP (pH 7.25). Tetrodotoxin (TTX, 0.5 µM) and gabazine (10 µM) were added to the ASCF to block voltage-gated Na^+^ channels and GABA_A_ receptors, respectively. For recording eEPSCs, whole-cell voltage-clamp measurements were performed at a holding potential of −70 mV. Schaffer collaterals were electrically stimulated using an ACSF-filled glass pipette placed in *stratum radiatum* (resistance about 1 MΩ) and a Model 2100 isolated pulse stimulator (A-M Systems). After finding a stimulation site, cells were allowed to recover for five minutes before the start of recording. Per condition, each cell was stimulated 30 times in a paired-pulse manner (20 Hz) at an inter-trial interval of 10 s. Recording pipettes were filled with solution containing (in mM): 12 KCl, 123 K-gluconate, 5 NaCl, 0.2 EGTA, 10 HEPES, 1.8 Mg-ATP, 0.3 Na-GTP, 0.1 spermine, 10 phosphocreatine (pH 7.3). Gabazine (10 µM) was added to the ASCF to isolate eEPSCs. Only recordings with an access resistance below 30 MΩ were accepted. Series resistance compensation was not applied.

### Surgical preparation, anesthesia and animal monitoring for in vivo LFP recordings

For analgesia, a subcutaneous injection of 200 mg/kg metamizol (Novacen) was administered 30 minutes prior to the start of the preparation. Animals were then placed onto a warm platform and anesthetized with isoflurane (3.5% for induction, 1–2% for maintenance) in pure oxygen (flow rate: 1 l/min). The skin overlying the skull was disinfected and locally infiltrated with 2% lidocaine (s.c.) for local analgesia. Scalp and periosteum were removed, and a custom-made plastic chamber with a central borehole (Ø 3 mm) was fixed on the skull using cyanoacrylate glue (UHU) (P3–5: 3.5 mm rostral from lambda and 1.5 mm lateral from midline; P10–12: 3.5 mm rostral from lambda and 2 mm lateral from midline). A subcutaneous catheter was placed at the lateral abdomen to allow for an acute injection of C21 (3 mM dissolved in 0.9% NaCl) and vehicle, respectively. To this end, a small tube (outer diameter: 0.61 mm, inner diameter: 0.28 mm, length: 70 mm), prefilled with 0.9% NaCl solution (volume: 4 µl) was inserted with a guiding canula (19G) and fixed using cyanoacrylate glue. For the hippocampal window preparation (Graf et al., 2021), the plastic chamber was tightly connected to a preparation stage and subsequently perfused with warm artificial cerebrospinal fluid (ACSF) containing (in mM): 125 NaCl, 4 KCl, 25 NaHCO_3_, 1.25 NaH_2_PO_4_, 2 CaCl_2_, 1 MgCl_2_ and 10 glucose (pH 7.4, 35–36°C). A circular hole was drilled into the skull using a tissue punch (outer diameter 2.7 mm). The underlying cortical tissue and parts of corpus callosum were carefully removed by aspiration using a vacuum supply and a blunt 27G or 30G needle. Care was taken not to damage alveus fibers. As soon as bleeding stopped, the animal was transferred to the microscope stage.

During *in vivo* recordings, body temperature was continuously monitored and maintained at close to physiological values (36–37°C) by means of a heating pad and a temperature sensor placed below the animal. Spontaneous respiration was monitored using a differential pressure amplifier (Spirometer Pod and PowerLab 4/35, ADInstruments). Isoflurane was discontinued after completion of the surgical preparation and gradually substituted with the analgesic-sedative nitrous oxide (up to the fixed final N_2_O/O_2_ ratio of 3:1, flow rate: 1 l/min). Experiments commenced 60 min after withdrawal of isoflurane. At the end of the experiment, the animal was decapitated under deep isoflurane anesthesia.

### In vivo LFP recordings

The recording chamber was continuously perfused with warm ACSF (as above). A tungsten microelectrode (Tunglass-1, 0.8 MΩ impedance, Kation Scientific) was lowered just above the hippocampal formation. ACSF was then removed and the hippocampal window was filled up with agar (1.5%, in 0.9 mM NaCl) and covered with a custom-made cover glass that allowed positioning of the electrode. As soon as the agar solidified, the chamber was reperfused with ACSF. Before inserting the electrode into the brain, an LFP reference signal was recorded for 5 minutes to estimate the noise picked up by the electrode (technical noise). Next, the microelectrode was slowly advanced into *stratum radiatum* (200–250 µm below the hippocampal surface). The final electrode depth was determined when the recorded eSPWs showed a polarity reversal. Electrophysiological signals were acquired and band-pass-filtered at 3–3,000 Hz using an EXT-02F/2 amplifier (npi electronic), a 16-bit AD/DA board (PowerLab 4/35, ADInstruments) and the software LabChart 8 (ADInstruments). Signals were sampled at 20 kHz. After recording spontaneous activity for 30 min, C21 was injected (s.c.) via the catheter at a concentration of 3 mg/kg. NaCl (0.9%) served as the vehicle control. The LFP recording continued for at least 2 hours after injection of C21.

### Analysis and quantification

#### Two-photon Ca^2+^ imaging in vitro

Image stacks were registered using NoRMCorre (Pnevmatikakis and Giovannucci, 2017). Time periods with residual z-drift were visually identified and considered as missing values in all subsequent analyses. Regions of interest (ROIs) corresponding to the somata of putative CA1 PCs were manually selected (Fiji). Subsequent analyses were performed using custom scripts in Matlab. For each cell, the time-course of fluorescence *F*(*t*) was obtained by framewise averaging across all pixels belonging to its ROI. Baseline fluorescence *F*_0_(*t*) was computed by oversmoothing *F*(*t*) using a second-order Savitzky-Golay algorithm (window length, 1,500 frames). Fluorescence signals were then expressed as relative changes from baseline levels, i.e. Δ*F*⁄*F*_0_ (*t*). Ca^2+^ transients (CaTs) were detected from Δ*F*⁄*F*_0_ (*t*) traces using UFARSA (Rahmati et al., 2018). Network events were operationally defined as GDPs by following a previous approach (Flossmann et al., 2019): (1) CaTs were classified as GDP-related if they fell into a 500-ms-long time interval during which the fraction of active cells (i.e., cells with ≥1 detected CaT) was ≥20%. (2) Neighboring GDP-related CaTs were assigned to the same GDP if both shared a 500-ms-long time interval during which the fraction of active cells was ≥20%, otherwise to separate GDPs.

#### Electrophysiological recordings in vitro

Resting membrane potential was determined as the median of the recorded voltage *V*_*m*_(*t*) (at zero current) in consecutive 20-s intervals. Membrane resistance was estimated under current-clamp conditions from the steady-state voltage change evoked by 500-ms-long hyperpolarizing test pulses (−10 pA at P2–5, −20 pA at P8–12, applied at 20-s intervals). For AP detection, we computed the first derivative of *V*_*m*_(*t*) and smoothed it using a second-order Savitzky-Golay algorithm (window: 5 samples), yielding 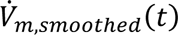. Time points at which 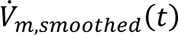 crossed a threshold of 20 V/s were considered as AP onsets. The value of *V*_*m*_(*t*) corresponding to the onset of the first AP evoked at the lowest amplitude of injected current was defined as AP threshold. The maximum mean AP frequency was computed as the maximum number of evoked APs per current step divided by step duration. The maximum instantaneous AP frequency was determined as the inverse of the minimum interval between two successive evoked APs. Rheobase was estimated as the minimum amplitude of injected current that gave rise to at least one AP. The quantities were computed separately for each of the three series of current injections per condition. For statistics, we computed the mean across the three series, except for rheobase in which case the median was used. Analyses were performed using custom scripts in Matlab. mEPSCs were detected using Minianalysis 6.0 (Synaptosoft). Events were visually selected and semi-automatically detected based on an amplitude and area criterion. eEPSCs were analyzed using Clampfit 11.2 (pClamp 11.2, Molecular Devices). eEPSCs were visually classified as failures and successes, respectively. To correct for a minor contamination of eEPSCs by the late decay of the stimulation artefact, eEPSC amplitudes were computed by subtracting the mean amplitude of failures from raw eEPSC amplitudes.

#### In vivo LFP recordings

The LFP was analyzed using custom-written Matlab scripts. Artefacts and noise peaks were removed semi-automatically. The LFP signal was downsampled to 500 Hz and baseline-corrected using a median filter (window length: 1 s). As an unbiased approach to quantify the input to CA1, we calculated the Welch’s power spectral density estimate (window length: 6 s, window overlap: 20%) for a 30-min window during control and at 30–60 min after C21 injection. The LFP reference signal was used for normalization. At P3–5, eSPWs were detected based on amplitude (larger than −0.05 mV) and an increase in bandpower (15–30 Hz). Continuous spectrograms were computed using the toolbox chronux (version 2.12, http://chronux.org/) (window length: 1 s, number of tapers: 3, window overlap: 80%; frequency range: 3–100 Hz, resulting in a datapoint every 0.2 s).

### Statistical analysis

Statistical analyses were performed using Matlab (2021b), OriginPro (2020) and IBM SPSS Statistics (28). Population data are reported as mean ± standard error of the mean (SEM), unless stated otherwise. Biological replicates (n) are given in Results and Extended Data Table 1-1. For significance testing, multi-group data were fitted with generalized linear mixed models (GLMM). The probability distribution of the dependent variable was specified as normal (with identity link) or gamma (with log link) based on *a-priori* considerations and the visual evaluation of the goodness of fit. We defined ‘genotype’ (or ‘dataset’), ‘condition’ and their interaction as fixed effects factors. We used ‘condition’ to identify repeated observations on the same subject and ‘subject’ as random effects. If the corrected (overall) model was non-significant, fixed effects were not further evaluated. If the corrected (overall) model and the interaction term were significant, we additionally computed the simple contrasts (P values adjusted for multiple comparisons using the Holm-Bonferroni method). In the figures, significance indicators refer to the P values of simple contrasts. P values (two-tailed tests) lower than 0.05 were considered statistically significant. Detailed statistical information is provided in Extended Data Table 1-1.

